# Genomic basis of rapid urban evolution revealed by the subgenome-resolved genome of octoploid *Oxalis corniculata*

**DOI:** 10.64898/2026.04.21.719839

**Authors:** Hideaki Iimura, Mitsuhiko P. Sato, Yuta B. Aoyagi, Shinji Kikuchi, Yuuya Tachiki, Kei Uchida, Koki Katsuhara, Kiriko Hiraoka, Yuya Fukano, Kenta Shirasawa

## Abstract

Urbanization is a major driver of contemporary evolution, yet the genomic basis of urban adaptation remains poorly understood, particularly in non-model plants with complex polyploid genomes. Here, we investigate the genetic mechanisms underlying leaf color variation in the octoploid *Oxalis corniculata*, a phenotype associated with heat tolerance in urban environments. By integrating high-fidelity long-read sequencing and chromosome conformation capture, we generated the subgenome-resolved, chromosome-scale genome assemblies for both red- and green-leaved lines, resolving four distinct subgenomes. LTR insertion timing revealed a two-step hybridization history that established this octoploid genome within the last 1 million years. Leveraging a nationwide citizen science initiative, we collected and analyzed over 1,700 samples across a broad geographic range. We identified a major locus on one subgenome underlying this variation and implicate a coding-sequence repeat-length polymorphism in a *MYB* transcription factor as the candidate causal variant. This simple sequence repeat likely acts as a molecular "tuning knob" for rapid adaptation to urban heat islands. This study provides a new baseline for evolutionary ecological genomics in plants and highlights the power of integrating advanced genomics with public participation to forecast evolutionary responses in an increasingly urbanized world.

## Introduction

Urbanization is one of the fastest and most widespread forms of environmental change on Earth (Seto et al., 2011; Seto et al., 2012). Cities impose strong and often repeatable selection pressures through elevated temperatures, altered water availability, artificial light, pollution, and chronic disturbance, making urban landscapes powerful natural experiments for studying rapid evolution (Johnson and Munshi-South, 2017). Urban ecosystems also provide model systems for empirically testing eco-evolutionary interactions, as they allow us to examine how changes in landscape structure alter selection and connectivity, thereby promoting evolutionary change (Fukano et al., 2023a). Since the classic example of industrial melanism, numerous studies have reported repeated phenotypic divergence between urban and non-urban populations across diverse taxa (Kettlewell, 1955; Alberti *et al*., 2017; Merckx *et al*., 2021; Capilla-Lasheras *et al*., 2022; Sandmeyer *et al*., 2025). However, remarkably few have identified the specific selective agents driving these differences and the genomic basis underlying them (Van’t Hof *et al*., 2016; Lambert *et al*., 2021; Santangelo *et al*., 2022; Winchell *et al*., 2023).

This gap is particularly acute in non-model plants, where large and frequently polyploid genomes, together with limited genomic resources, make it difficult to infer the genetic basis of urban adaptation and the eco-evolutionary dynamics of populations (Phillips, 2024; Karbstein *et al*., 2025). As a result, the repeatability of urban evolution, the genomic consequences of urban landscape changes, and the historical processes by which adaptive variants arise and spread remain poorly resolved (Lambert *et al*., 2021). Bridging this gap requires intensive eco-evolutionary genomic approaches, including chromosome-scale reference genomes, population-scale sampling across many cities, spatially explicit population genomics, and the genome-wide mapping of adaptive loci.

*Oxalis corniculata* (creeping woodsorrel) serves as an ideal model system for addressing these challenges. A member of the genus *Oxalis* distributed as a cosmopolitan weed, *O. corniculata* is characterized by three heart-shaped leaflets—a morphology so well-conserved within the genus that it is frequently misidentified as *O. debilis*, *O. dillenii*, or *O. pes-caprae*, and sometimes confused with clovers (*Trifolium* spp.). In Japan, *Oxalis* has been culturally significant since the Heian period (794–1185), often used in family crests (kamon), and its heritable red- and green-leaf phenotypic variation was documented as early as 1709 in Edo-period herbalism texts (Kaibara, 1709). Our previous research demonstrated that these phenotypes exhibit a striking distribution: green-leaved plants dominate rural habitats, while red-leaved individuals are disproportionately common in urban environments (Fukano et al., 2023b). Experimental evidence further indicates that red leaves, characterized by anthocyanin accumulation, confer higher heat tolerance, suggesting that the urban heat island effect acts as a key selective agent favoring this trait.

Identifying the genetic basis of this repeatable urban evolution, however, presents major challenges. While anthocyanin biosynthesis is a well-studied pathway, the specific molecular mechanisms driving urban adaptation in *O. corniculata* remain unclear. Furthermore, because urban environments have established independently worldwide, it remains unknown whether the underlying molecular evolution is shared across cities or represents independent events. Resolving this requires both high-quality genomic resources and extensive, spatially replicated sampling. Although genome sequences have been reported for some relatives, such as the diploid *O. articulata* (Yang et al., 2025), *O. oulophora* (Vanrooyen and Emshwiller, 2026), and the allotetraploid *O. stricta* (Wood et al., 2025), no reference genome for *O. corniculata* is currently available. This is largely due to its complex genomic architecture, characterized by a ploidy series (including tetraploids and octoploids) with a base chromosome number of 6 (Marks, 1956).

We addressed these difficulties by launching the Minna de Katabami ("Oxalis, Together") Project (https://sites.google.com/kazusa.or.jp/oxalis/), an open genomics initiative collaborating with citizen scientists. Building on previous efforts that collected samples from limited locations (Shirasawa et al., 2021; Fukano et al., 2023b; Shirasawa et al., 2024), we leveraged a nationwide network to achieve unprecedented sampling density across the Japanese archipelago. Simultaneously, we utilized advanced sequencing technologies, including PacBio high-fidelity (HiFi) long-reads and high-throughput chromosome conformation capture (Dudchenko et al., 2017), to overcome the hurdles of polyploid assembly.

In this study, we provide the subgenome-resolved near-telomere-to-telomere genome sequences for both red- and green-leaved octoploid *O. corniculata* (Figure 1). Using these resources, combined with SNP genotyping analysis of the citizen-collected samples, we identify the causal locus and a candidate variant (an AAT simple sequence repeat) underlying the red-leaf phenotype. Our results clarify the population structure and demonstrate the repeatability of urban adaptation, providing a new baseline for evolutionary ecological genomics in non-model organisms.

**Figure 1.**
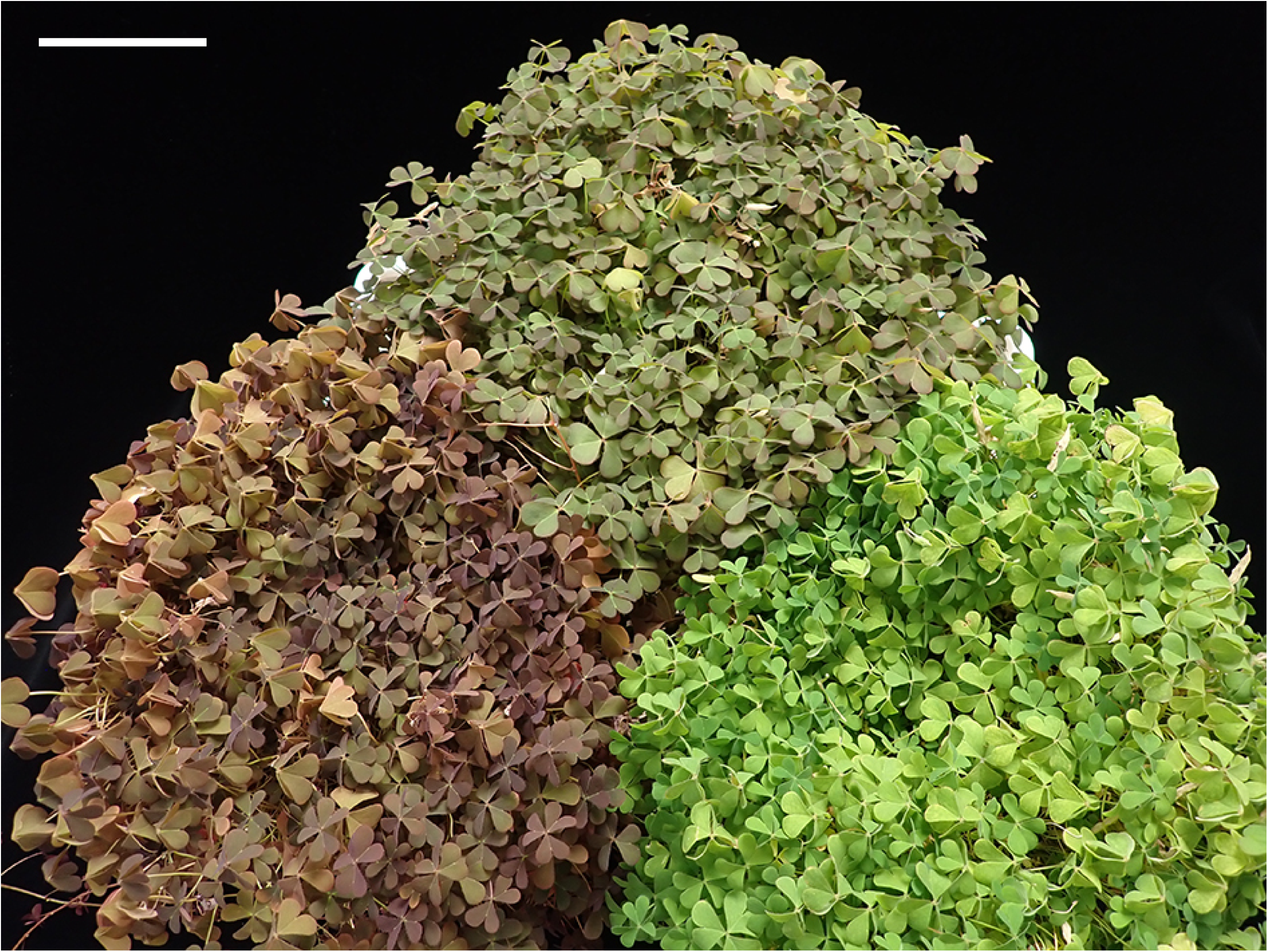
Aerial parts of *Oxalis corniculata*. Representative images of the red-leaved line (bottom left), the green-leaved line (bottom right), and their F1 hybrid (top center). Scale bar = 10 cm.

## Results

### Chromosome observation and genome size estimation

Prior to genome sequencing and analysis, the chromosome numbers of the red- and green-leaved lines of *O. corniculata* were investigated. A total of 48 chromosomes were observed in metaphase cells of buds from both lines (Supplementary Figure S1), consistent with a previous report (Marks, 1956). This result indicates that both the red-and green-leaved lines are octoploid, with a basic chromosome number of six (2n = 8x = 48).

To estimate the genome sizes of the red- and green-leaved lines, a total of 113.2 Gb and 115.3 Gb of short-read data, respectively, were subjected to *k*-mer distribution analysis. The results showed two prominent peaks alongside two minor peaks (Supplementary Figure S2), further suggesting that the red- and green-leaved lines possess octoploid genomes. The haploid genome size of *O. corniculata* was estimated to be ranging from approximately 285 Mb to 306 Mb.

### Genome sequencing and chromosome-scale assembly

A total of 28.8 Gb of HiFi reads and 411.7 million (M) Omni-C paired-end reads were obtained from the red-leaved line. These reads were assembled into 368 contigs with an N50 of 38.8 Mb and a total length of 1,016.4 Mb. After removing potential haplotigs and overlap sequences, the resultant contig size was 959 Mb consisting of 41 contig sequences. (Supplementary Table S1). In parallel, a total of 29.0 Gb of HiFi reads and 406.1 M Omni-C paired-end reads from the green-leaved line were assembled into 159 contigs with an N50 of 40.0 Mb and a total length of 967.6 Mb. Similarly, potential haplotigs and overlapping sequences were purged, yielding an assembly size of 954 Mb consisting of 29 contig sequences (Supplementary Table S1).

The contig sequences of the red- and green-leaved lines were aligned. In total, 41 contigs from the red-leaved line and 25 contigs from the green-leaved line were complementarily aligned to construct 24 pseudomolecule sequences (Table 1). The remaining four short contigs (534 kb in total) from the green-leaved line were not integrated into the pseudomolecules but were retained as unplaced contigs in the genome assembly. The pseudomolecule sequences of the red- and green-leaved lines were designated OCOR_r3.0 and OCOG_r3.0, respectively.

**Table 1.**
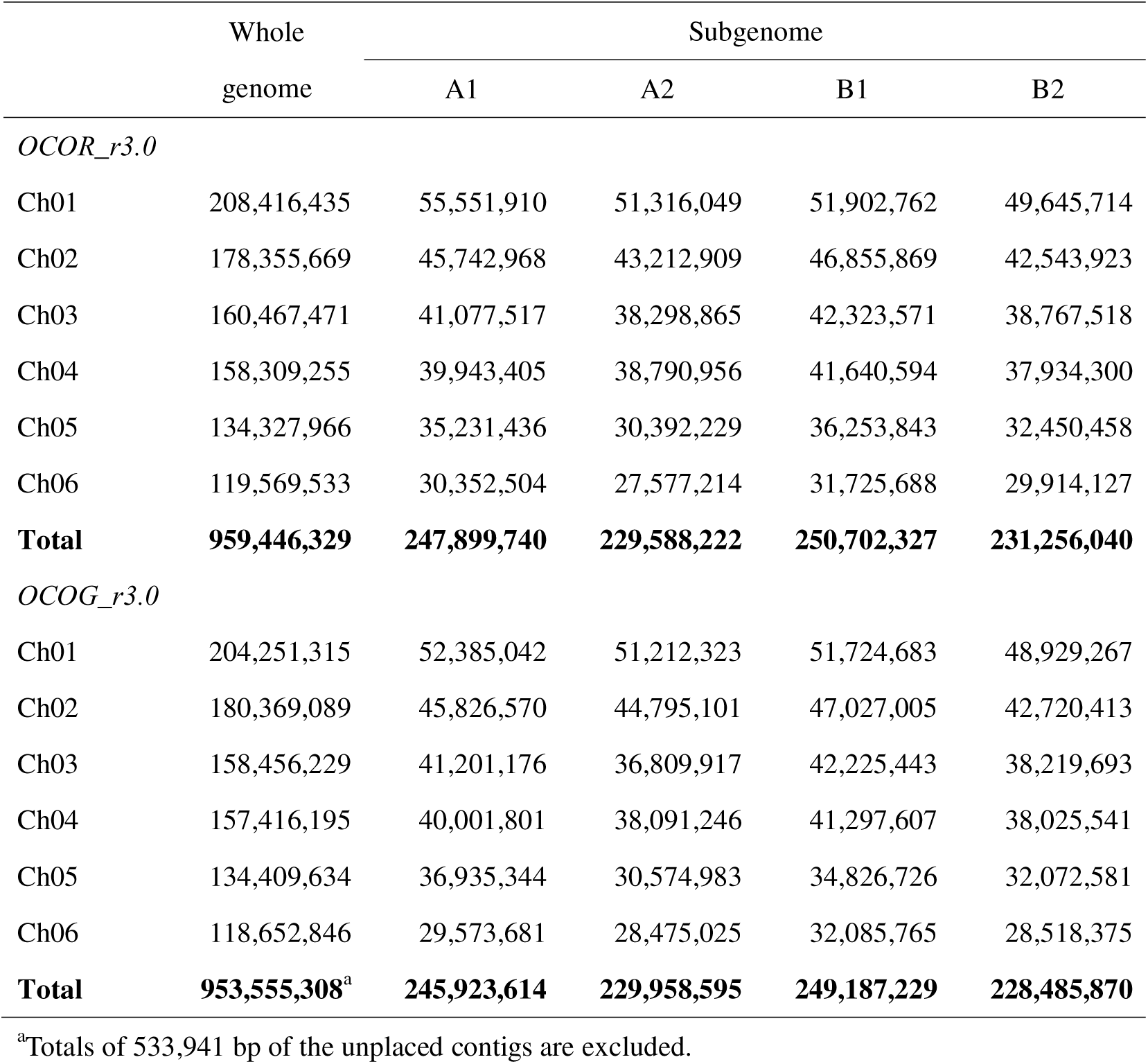
Assembly statistics for the haplotype-phased, chromosome-level genomes of *Oxalis corniculata*.

|  | Whole | Subgenome |  |  |  |
| --- | --- | --- | --- | --- | --- |
|  | genome | A1 | A2 | B1 | B2 |
| *OCOR_r3.0* |  |  |  |  |  |
| Ch01 | 208,416,435 | 55,551,910 | 51,316,049 | 51,902,762 | 49,645,714 |
| Ch02 | 178,355,669 | 45,742,968 | 43,212,909 | 46,855,869 | 42,543,923 |
| Ch03 | 160,467,471 | 41,077,517 | 38,298,865 | 42,323,571 | 38,767,518 |
| Ch04 | 158,309,255 | 39,943,405 | 38,790,956 | 41,640,594 | 37,934,300 |
| Ch05 | 134,327,966 | 35,231,436 | 30,392,229 | 36,253,843 | 32,450,458 |
| Ch06 | 119,569,533 | 30,352,504 | 27,577,214 | 31,725,688 | 29,914,127 |
| **Total** | **959,446,329** | **247,899,740** | **229,588,222** | **250,702,327** | **231,256,040** |
| *OCOG_r3.0* |  |  |  |  |  |
| Ch01 | 204,251,315 | 52,385,042 | 51,212,323 | 51,724,683 | 48,929,267 |
| Ch02 | 180,369,089 | 45,826,570 | 44,795,101 | 47,027,005 | 42,720,413 |
| Ch03 | 158,456,229 | 41,201,176 | 36,809,917 | 42,225,443 | 38,219,693 |
| Ch04 | 157,416,195 | 40,001,801 | 38,091,246 | 41,297,607 | 38,025,541 |
| Ch05 | 134,409,634 | 36,935,344 | 30,574,983 | 34,826,726 | 32,072,581 |
| Ch06 | 118,652,846 | 29,573,681 | 28,475,025 | 32,085,765 | 28,518,375 |
| **Total** | **953,555,308<sup>a</sup>** | **245,923,614** | **229,958,595** | **249,187,229** | **228,485,870** |

<sup>a</sup>Totals of 533,941 bp of the unplaced contigs are excluded.

Complete BUSCO scores for OCOR_r3.0 and OCOG_r3.0 were both 98.7% (Table 2). In OCOR_r3.0, telomere repeats were found at both ends of 22 sequences and at one end of the remaining two sequences. In OCOG_r3.0, telomere repeats were found at both ends of 21 sequences and at one end of the remaining three sequences.

**Table 2.** Assessment of genome assembly completeness using BUSCO for the red- and green-leaved lines.

|  | Whole genome |  |  | Subgenome |  |  |  |
| --- | --- | --- | --- | --- | --- | --- | --- |
|  | A1+A2+B1+B2 | A1+A2 | B1+B2 | A1 | A2 | B1 | B2 |
| *OCOR_r3.0* |  |  |  |  |  |  |  |
| Complete | 99.0 | 84.9 | 90.3 | 74.2 | 74.4 | 80.3 | 80.7 |
| Single-copy complete | 7.4 | 25.6 | 23.8 | 71.9 | 70.9 | 77.3 | 76.0 |
| Duplicated complete | 91.6 | 59.3 | 66.5 | 2.3 | 3.5 | 3.0 | 4.7 |
| Fragmented | 0.6 | 7.7 | 6.0 | 13.5 | 10.2 | 11.1 | 9.5 |
| Missing | 0.4 | 7.4 | 3.7 | 12.3 | 15.4 | 8.6 | 9.8 |
| *OCOG_r3.0* |  |  |  |  |  |  |  |
| Complete | 99.1 | 85.2 | 90.4 | 73.3 | 74.5 | 80.4 | 81.0 |
| Single-copy complete | 8.0 | 27.0 | 23.9 | 71.1 | 71.3 | 77.4 | 76.5 |
| Duplicated complete | 91.1 | 58.2 | 66.5 | 2.2 | 3.2 | 3.0 | 4.5 |
| Fragmented | 0.6 | 6.8 | 5.6 | 13.0 | 9.4 | 10.2 | 9.5 |
| Missing | 0.3 | 8.0 | 4.0 | 13.7 | 16.1 | 9.4 | 9.5 |

### Haplotype-phased subgenome identification and estimation of divergence times

The pseudomolecule sequences of OCOR_r3.0 and OCOG_r3.0 were compared (Figure 2). Based on sequence similarity, the 24 pseudomolecules for each line were divided into two major groups (A and B), each consisting of two subgroups. Each subgroup contained six sequences.

**Figure 2.**
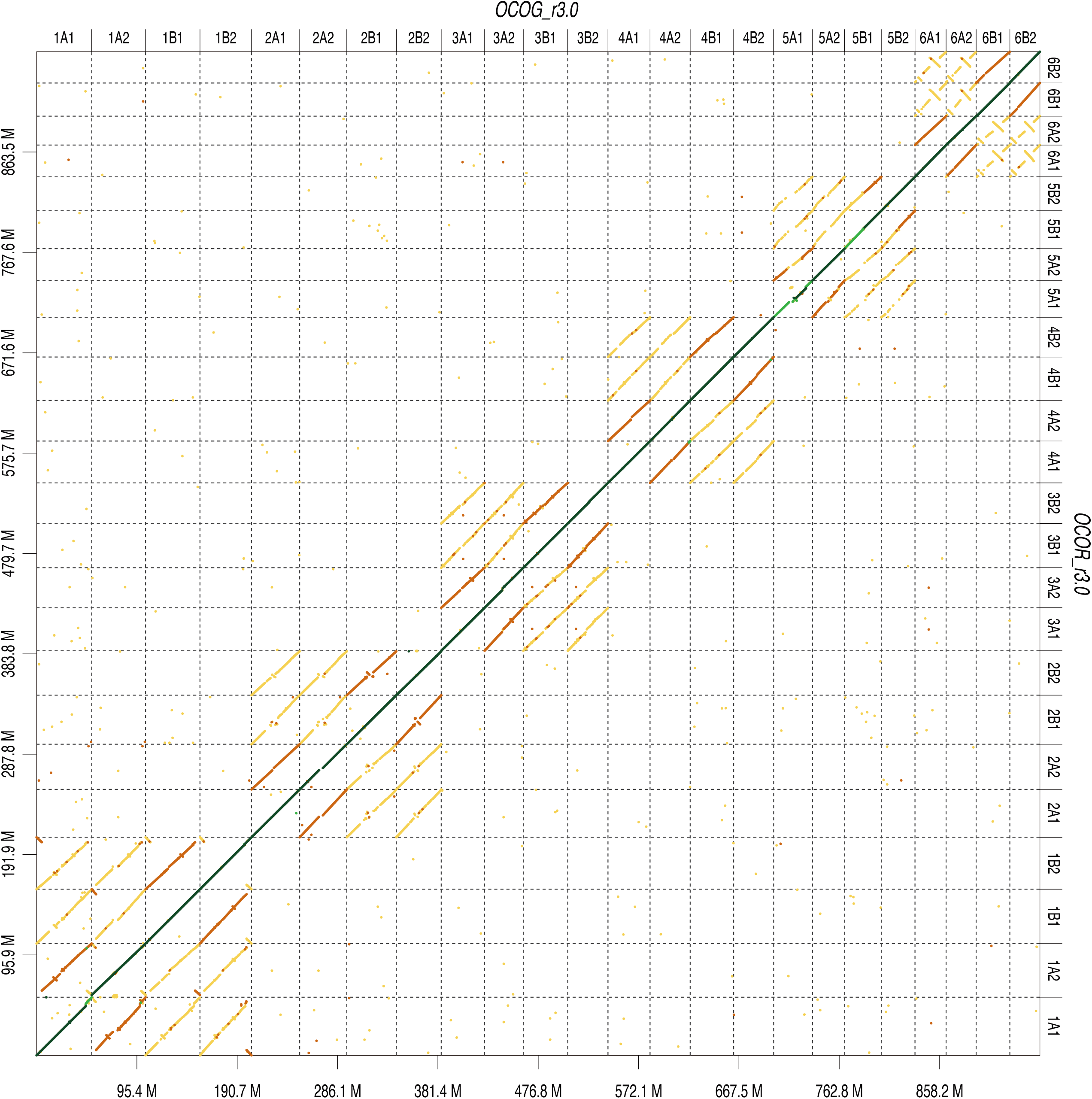
Genome synteny between the red- and green-leaved lines of *Oxalis corniculata*. A dot plot comparison of the chromosome-scale sequences. The x-axis represents the green-leaved line assembly (OCOG_r3.0), and the y-axis represents the red-leaved line assembly (OCOR_r3.0).

We then analyzed the repetitive sequences in the four subgroups. Repeat sequences occupied 430.9 Mb (44.9%) and 443.2 Mb (46.5%) of the red- and green-leaved *O. corniculata* genomes, respectively (Table 3). Long terminal repeat (LTR) elements were the most abundant repetitive sequences (14.6% in OCOR_r3.0 and 14.2% in OCOG_r3.0), followed by DNA transposons (5.6% in OCOR_r3.0 and 5.3% in OCOG_r3.0). Unclassified repeats constituted 18.8% and 20.1% of the red- and green-leaved *O. corniculata* genomes, respectively. Among the classified repeats, 32 types were enriched in group A and depleted in group B (Supplementary Figure S3A), whereas 24 types were depleted in group A and enriched in group B (Supplementary Figure S3B). We therefore defined groups A and B as subgenomes A and B, respectively. Furthermore, 40 repeat types were enriched in one subgroup of subgenome A and one subgroup of subgenome B, but depleted in the remaining subgroups (Supplementary Figure S3C). Consequently, we defined the repeat-rich subgroups as A1 and B1, and the repeat-poor subgroups as A2 and B2. To validate these definitions, repetitive *k*-mers serving as "differential signatures" (Jia *et al*., 2022) within the four subgenome sequences were compared to phase the subgenomes. Subgenome-specific regions and LTR retrotransposons were detected across the four subgenomes, and as expected, the subgenome classifications defined above were fully supported by this k-mer analysis (Supplementary Figure S4A-D).

**Table 3.** Classification and proportion of repetitive elements in the red- and green-leaved *Oxalis corniculata* genomes.

| Repeat type | OCOR_r3.0 |  |  | OCOG_r3.0 |  |  |
| --- | --- | --- | --- | --- | --- | --- |
|  | No. of repeats | Total length (bp) | Relative proportion (%) | No. of repeats | Total length (bp) | Relative proportion (%) |
| SINEs | 10,006 | 1,598,962 | 0.2 | 9,197 | 1,572,192 | 0.2 |
| LINEs | 66,230 | 32,164,370 | 3.4 | 54,304 | 32,546,344 | 3.4 |
| LTR elements | 174,170 | 139,936,659 | 14.6 | 177,532 | 135,811,847 | 14.2 |
| DNA transposons | 171,604 | 54,047,678 | 5.6 | 153,613 | 50,656,256 | 5.3 |
| Small RNA | 5,551 | 1,989,711 | 0.2 | 6,202 | 2,156,897 | 0.2 |
| Satellites | 37,271 | 4,254,873 | 0.4 | 20,671 | 10,608,353 | 1.1 |
| Simple repeats | 210,679 | 7,947,803 | 0.8 | 210,517 | 7,955,123 | 0.8 |
| Low complexity | 35,935 | 1,669,032 | 0.2 | 36,605 | 1,700,835 | 0.2 |
| Unclassified | 708,776 | 179,868,483 | 18.8 | 702,623 | 192,025,986 | 20.1 |

In the red-leaved line, the insertion times of subgenome-specific LTRs were estimated at 4.1 and 5.3 million years ago (MYA) for subgenomes A (A1 and A2) and B (B1 and B2), respectively (Supplementary Figure S4E), and at 0.7 and 0.5 MYA for subgenomes 1 (A1 and B1) and 2 (A2 and B2) (Supplementary Figure S4F). Similarly, in the green-leaved line, these insertion times were estimated at 4.8 and 5.4 MYA for subgenomes A and B (Supplementary Figure S4G), and 0.7 and 0.4 MYA for subgenomes 1 and 2 (Supplementary Figure S4H).

### Protein-coding gene prediction

To predict genes within the genome assemblies, full-length cDNA sequences were obtained from the red- and green-leaved lines. A total of 83,786 and 83,722 non-redundant transcript sequences for the red- and green-leaved lines, respectively, were used for gene prediction.

In OCOR_r3.0, a total of 117,631 genes were predicted (Table 4), of which 57,378 and 60,253 genes were located in subgenomes A and B, respectively. The numbers of predicted genes in subgenomes A1, A2, B1, and B2 were 28,763, 28,615, 29,914, and 30,339, respectively. The complete BUSCO score for OCOR_r3.0 was 99.0%, while scores for subgenomes A and B were 84.9% and 90.3%, respectively. Complete BUSCO scores for subgenomes A1, A2, B1, and B2 were 74.2%, 74.4%, 80.3%, and 80.7%, respectively.

**Table 4.** Statistics of protein-coding genes predicted in the red- and green-leaved *Oxalis corniculata* assemblies.

|  | Whole genome | Subgenome |  |  |  |
| --- | --- | --- | --- | --- | --- |
|  |  | A1 | A2 | B1 | B2 |
| *OCOR_r3.0* |  |  |  |  |  |
| Ch01 | 26,236 | 6,468 | 6,353 | 6,583 | 6,832 |
| Ch02 | 22,957 | 5,740 | 5,682 | 5,745 | 5,790 |
| Ch03 | 20,443 | 4,996 | 4,939 | 5,227 | 5,281 |
| Ch04 | 19,757 | 4,801 | 4,907 | 4,948 | 5,101 |
| Ch05 | 13,066 | 3,061 | 3,017 | 3,521 | 3,467 |
| Ch06 | 15,172 | 3,697 | 3,717 | 3,890 | 3,868 |
| **Total** | **117,631** | **28,763** | **28,615** | **29,914** | **30,339** |
| *OCOG_r3.0* |  |  |  |  |  |
| Ch01 | 26,222 | 6,365 | 6,385 | 6,667 | 6,805 |
| Ch02 | 23,355 | 5,848 | 5,847 | 5,830 | 5,830 |
| Ch03 | 20,537 | 5,039 | 5,015 | 5,230 | 5,253 |
| Ch04 | 20,063 | 4,883 | 4,982 | 5,015 | 5,183 |
| Ch05 | 13,243 | 3,161 | 3,088 | 3,447 | 3,547 |
| Ch06 | 15,244 | 3,773 | 3,767 | 3,902 | 3,802 |
| **Total** | **118,744<sup>a</sup>** | **29,069** | **29,084** | **30,091** | **30,420** |

<sup>a</sup>Totals of 80 genes predicted on the unplaced contigs are excluded.

In OCOG_r3.0, a total of 118,744 genes were predicted (Table 4), of which 58,153 and 60,511 genes were located in subgenomes A and B, respectively, while the remaining 80 genes were found in the four unplaced contigs. The numbers of predicted genes in subgenomes A1, A2, B1, and B2 were 29,069, 29,084, 30,091, and 30,420, respectively. The complete BUSCO score for OCOG_r3.0 was 99.1%, with scores for subgenomes A and B reaching 85.2% and 90.4%, respectively. Complete BUSCO scores for subgenomes A1, A2, B1, and B2 were 73.3%, 74.5%, 80.4%, and 81.0%, respectively.

### Organelle genome assembly and gene annotation

The chloroplast genome assemblies for the red- and green-leaved lines were 152,165 bp and 152,164 bp in length, respectively (Figure S5A, B). Compared to the green-leaved line, the red-leaved line contained a single-base insertion at a poly-T site. In both the red- and green-leaved lines, the same 130 genes were predicted, including 84 protein-coding genes, 38 tRNA genes, and 8 rRNA genes. The mitochondrial genome assemblies for the red- and green-leaved lines, each consisting of two circular sequences with lengths of 196,770 bp and 225,435 bp, were completely identical (Figure S5C – F). The same 67 genes, including 39 protein-coding genes, 25 tRNA genes, and 3 rRNA genes, were predicted in the mitochondrial genomes of both lines.

### Genetic mapping of a gene for leaf color variation

To identify the genes responsible for leaf color variation, we crossed the red- and green-leaved lines used for genome sequencing to generate an F1 hybrid. The leaf color of the F1 plant was visually reddish-green (Figure 1); this was markedly different from the green-leaved parental line, but difficult to distinguish from the red-leaved parental line. The anthocyanin content index (ACI) for the red- and green-leaved parental lines was 10.5 and 3.6, respectively, while that for the F1 plant was 8.1. The F1 plant was then self-pollinated to generate an F2 population (n = 192). The number of red- (or reddish-green-) and green-leaved plants in the F2 population was 145 and 47, respectively. This segregation ratio suggested that red is semi-dominant over green and that leaf color is controlled by a single gene (P = 0.868, χ² = 0.028). The ACI for the F2 plants with red (or reddish-green) leaves ranged from 4.9 to 14.1, while those with green leaves ranged from 3.0 to 4.8 (Supplementary Figure S6A).

To identify the genetic locus controlling leaf color, we performed ddRAD-seq analysis. A mean of 4.95 M reads per sample (including F2, parental, and F1 lines) was obtained and mapped to OCOR_r3.0 with an alignment rate of 98.6%, leading to the detection of 575 SNPs. A genome-wide association study (GWAS) detected one significant peak for leaf color at 6,038,216 bp on chromosome 4B1 (Figure 3A). In parallel, sequence data were mapped to the OCOG_r3.0 reference genome with an alignment rate of 98.0%, detecting 548 SNPs. GWAS detected one significant peak at 5,869,588 bp on chromosome 4B1 (Supplementary Figure S7A). These two peaks were located in corresponding regions between the red- and green-leaved genomes.

**Figure 3.**
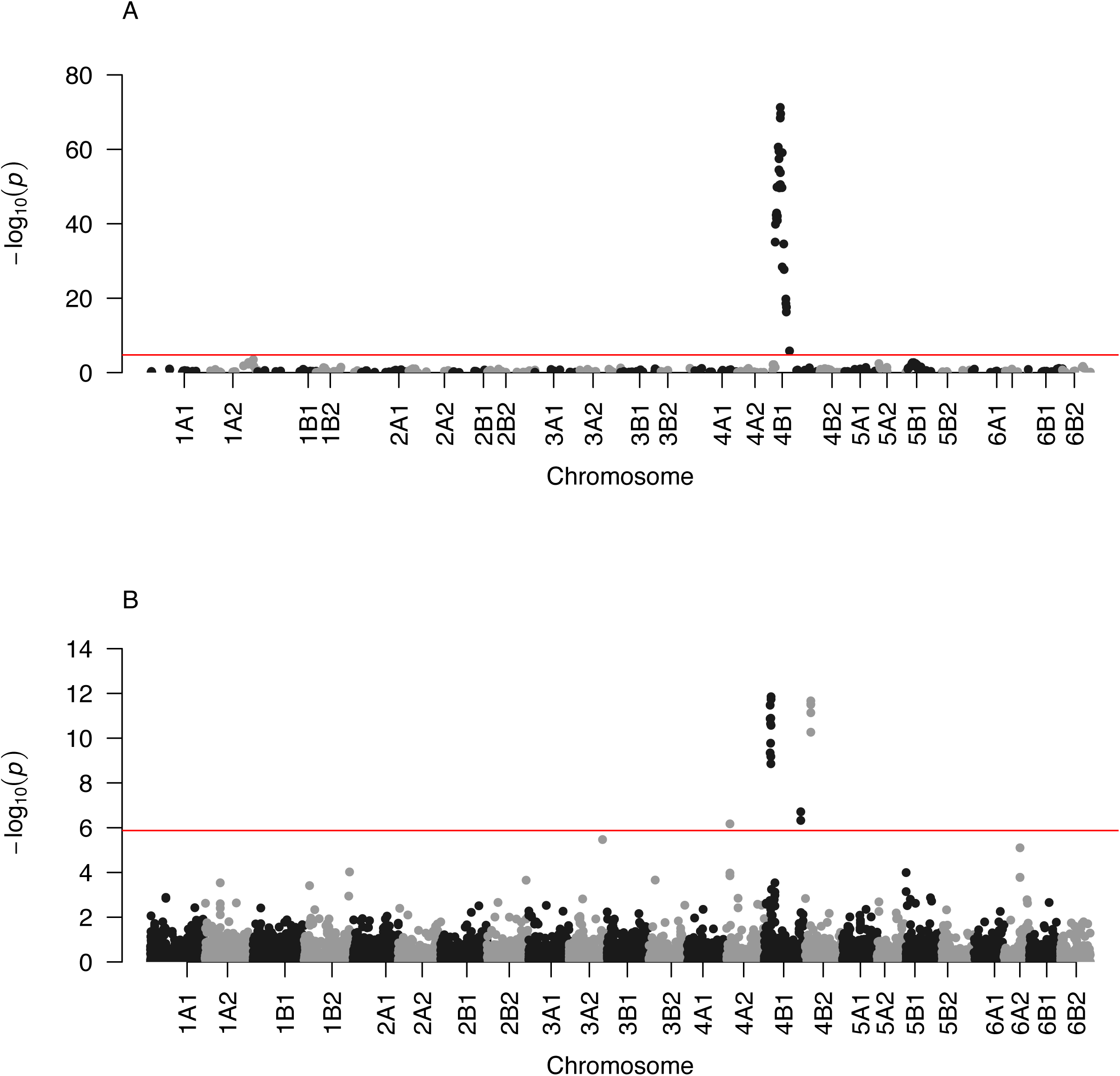
SNPs associated with leaf color variation identified by GWAS. Manhattan plots showing the distribution of SNPs associated with leaf color variation across the genome. The horizontal red lines indicate the Bonferroni-corrected significance threshold (α = 1%). Results are shown for (A) the F2 mapping population based on the red-leaved line genome as a reference and (B) the natural population collected from across Japan.

To narrow down the genomic region controlling leaf color, we analyzed the whole-genome SNP genotypes of 22 F2 plants that possessed chromosome recombination breakpoints in the regions flanking the GWAS peak. A mean of 153.1 M reads per sample was obtained and mapped to the reference genomes of the red-and green-leaved lines. The candidate region was narrowed to a 129-kb interval in OCOR_r3.0 and a 121-kb interval in OCOG_r3.0. Within this interval in OCOG_r3.0, we identified a gene encoding a partial MYB transcription factor. In accordance with the long-read transcriptome data, this gene consisted of three exons. However, no stop codon was found in the annotated exon sequences because the long-read sequencing prematurely terminated at a trinucleotide (AAT) repeat sequence within exon 3. To determine the full-length sequence of this MYB gene, we designed primer pairs and amplified cDNA spanning the AAT repeat. Sanger sequencing of the amplified cDNA revealed that exon 3 extended beyond the (AAT)_21_ repeat sequence and ended with a stop codon. The corresponding gene was also found in OCOR_r3.0. This allele also possessed the repetitive sequence, but it was 81 bases longer than the OCOG_r3.0 allele, configured as (AAT)_21_TGTTAT(AAT)_14_TGTTAT(AAT)_9_ (Figure 4).

**Figure 4.**
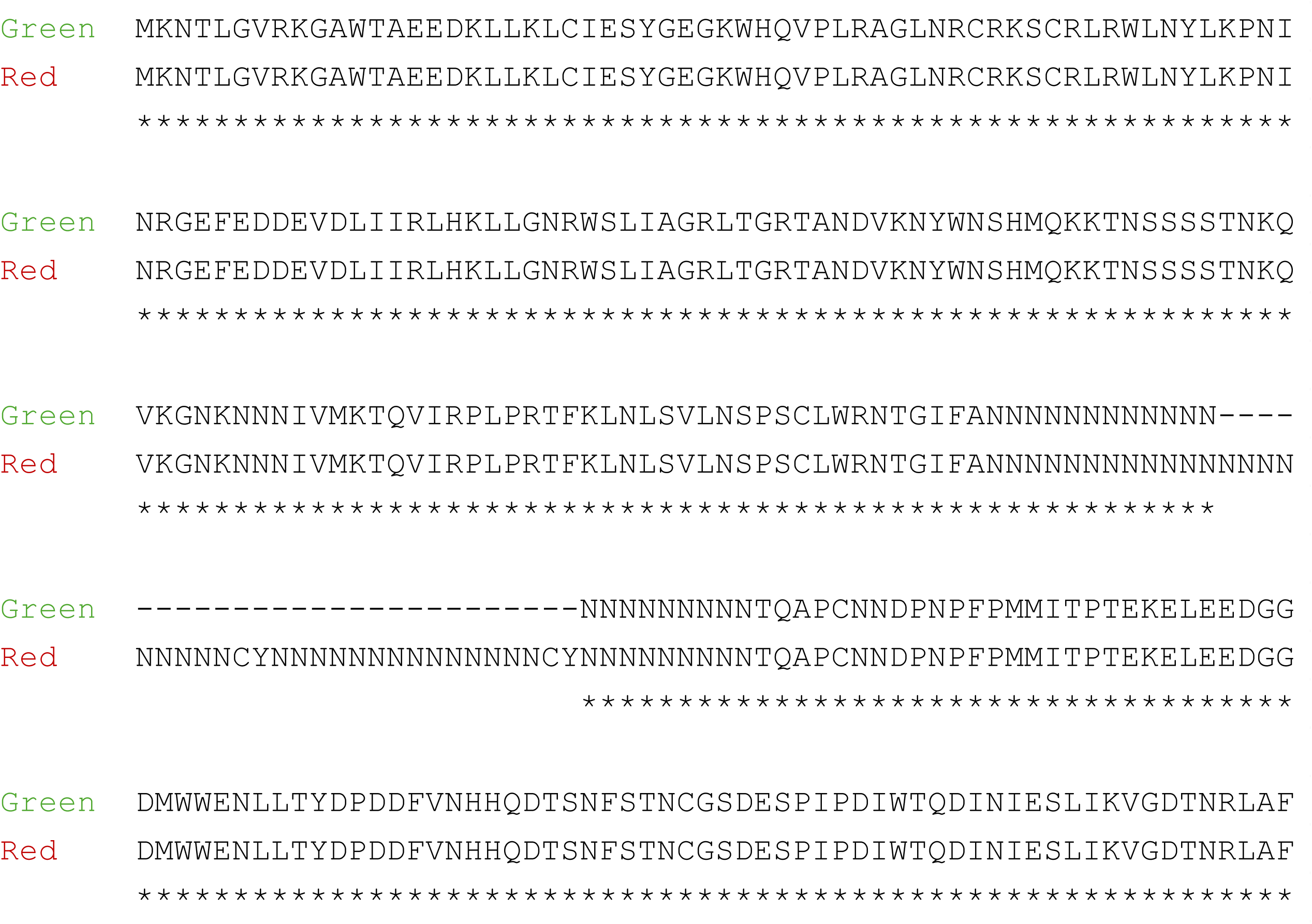
Multiple sequence alignment of the MYB protein underlying leaf color variation. Comparison of the amino acid sequences of the candidate gene between the red and green alleles.

### Genetic analysis of natural population

To investigate the distribution of this mutation in the field, we performed GWAS using 1,784 samples collected from 75 regions across Japan, which were contributed by participants of the Project (Supplementary Table S2). Because *O. corniculata* is morphologically similar to its relatives, we assumed the collected dataset might contain misidentified samples. Therefore, two *O. debilis*, four *O. dillenii*, and one *O. pes-caprae* sample were included as outgroup controls to filter out non-target species.

We performed ddRAD-seq analysis on all samples. A mean of 4.8 M raw reads per sample was obtained and mapped to OCOR_r3.0. This identified a total of 15,697 high-confidence SNPs. During quality control, 172 samples were excluded due to a high proportion of missing data, likely caused by low DNA quality or quantity (Supplementary Table S2). The remaining 1,612 samples, along with the seven control samples, were subjected to phylogenetic analysis. Of these, 323 samples clustered with the controls, suggesting they were either *O. debilis*, *O. dillenii*, or *O. pes-caprae* (Figure 5, Supplementary Table S2). The remaining 1,289 samples were confirmed as *O. corniculata*. Based on the phylogenetic analysis, these 1,289 samples were classified into three subpopulations (Figure 5, Supplementary Table S2): 208 samples in Group I, 456 samples in Group II, and 625 samples in Group III. The ACI values of the 1,289 samples ranged from 1.3 (green) to 14.8 (red), with an overall average of 4.6 (Supplementary Figure S6B). The mean ACI values for Groups I, II, and III were 4.0, 5.1, and 4.4, respectively. The red- and green-leaved lines used for reference genome sequencing belonged to Group III.

**Figure 5.**
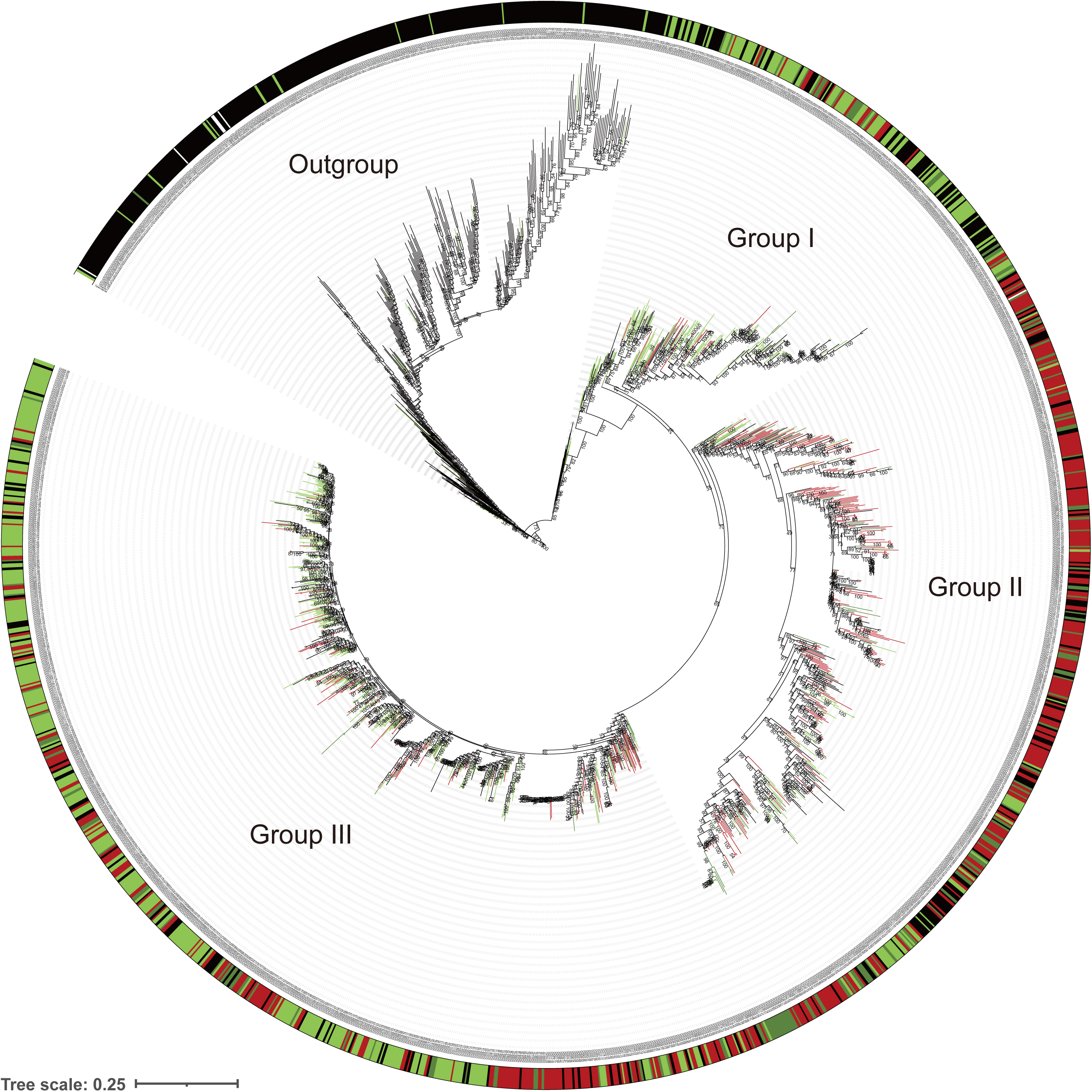
Phylogenetic relationship of *Oxalis corniculata* samples collected across Japan. A maximum-likelihood tree constructed using a genome-wide SNP dataset. The outer color bars indicate the phenotypic leaf color of each accession, as represented by the anthocyanin index (ACI).

To identify the genetic loci controlling leaf color across the natural population, GWAS was performed on the 1,289 *O. corniculata* samples. A prominent peak (P = 1.4 × 10^−12^) was detected at 6,948,984 bp on chromosome 4B1 (Figure 3B). When GWAS was performed independently for each subpopulation, significant peaks were detected in Group III (6,038,250 bp on chromosome 4B1; P = 1.6 × 10^−87^) and Group II (6,948,984 bp on chromosome 4B1; P = 7.0 × 10^−20^), but no significant peak was observed in Group I (Supplementary Figure S7B – D).

### Polymorphism analysis of the simple sequence repeat

To genotype the length variation of the repetitive sequence in the *MYB* gene, a primer pair was designed to amplify a 384-bp fragment from the red allele and a 303-bp fragment from the green allele. DNA fragments of the expected sizes were successfully amplified from the red- and green-leaved plants of the F2 population. Furthermore, within Group III of the natural population, the length of the repetitive sequence varied from 359 bp to 385 bp among red-leaved samples, and from 296 bp to 330 bp among green-leaved samples.

## Discussion

In this study, we integrated chromosome-scale genome sequences and large-scale population sampling enabled by citizen science. We mapped a major locus underlying red-leaf variation—a phenotype associated with heat tolerance in urban environments. These results provide a spatially explicit connection between landscape change and evolutionary change at the genomic level. Many studies of urban evolution rely on simple urban-versus-non-urban phenotypic contrasts or genome-wide scans (Thompson et al., 2016; Santangelo et al., 2018; Diamond and Martin, 2021; Santangelo et al., 2022). In contrast, we identified the candidate locus and allele underlying this phenotypic contrast, demonstrating that measurable landscape structure is associated with allele-frequency variation at this specific locus.

Several lines of evidence implicate a *MYB* gene at this locus as the strongest candidate for red-leaf variation. This specific region was supported by both F2 mapping and association analyses in natural populations (Figure 3, Supplementary Figure S7). The red and green alleles differ in the length and structure of an exonic AAT simple sequence repeat (SSR) (Figure 4), which is predicted to encode poly-asparagine (poly-N) peptides. Interestingly, this gene was initially missed by automated prediction and RNA-Seq mapping, likely because the repetitive (AAT)_n_ stretch disrupted sequence alignment. While a similar poly-N stretch exists in the MYB protein of *Rutidosis leptorrhynchoides* (XP_071724211), its functional role remains uncharacterized. MYB transcription factors are known regulators of anthocyanin biosynthesis (Naing and Kim, 2018; Yan et al., 2021). Most known variants involving *MYB* genes arise from cis-regulatory changes (Fattorini and Ó’Maoiléidigh, 2022); thus, the coding-sequence polymorphism identified here represents a potentially novel mechanism. From an evolutionary perspective, SSRs mutate faster than SNPs due to replication slippage (Hamblin et al., 2007), making them ideal "tuning knobs" for rapid adaptation in fluctuating urban environments (Horton et al., 2023; King, 2025).

Our high-quality genomic resources also provide insights into the evolutionary history of this octoploid species. We resolved four subgenomes (A1, A2, B1, and B2), each represented by six chromosomes (Table 1, Supplementaty Figure S3 and S4). Based on the distribution of subgenome-specific repetitive elements, we propose a two-step hybridization and polyploidization model to establish the octoploid genome of *O. corniculata* (Figure 6): Initially, two ancestral diploid lineages (A and B) diverged approximately 5 million years ago (MYA), as estimated by subgenome-specific LTR insertion times. These lineages hybridized to form an allotetraploid intermediate (AB). Subsequently, a second round of hybridization between differentiated tetraploid lineages (A1B1 and A2B2) occurred less than 1 MYA, resulting in the establishment of the extant octoploid *O. corniculata* (A1A2B1B2). Interestingly, BUSCO scores for individual subgenomes (73.3% - 81.0%) were lower than for the entire genome (99.0% and 99.1%), suggesting potential subfunctionalization or neofunctionalization following polyploidization. Despite this complexity, the genomic architecture and organelle sequences were highly conserved between red- and green-leaved lines (Figure 2), confirming that both lines indeed belong to *O. corniculata*.

**Figure 6.**
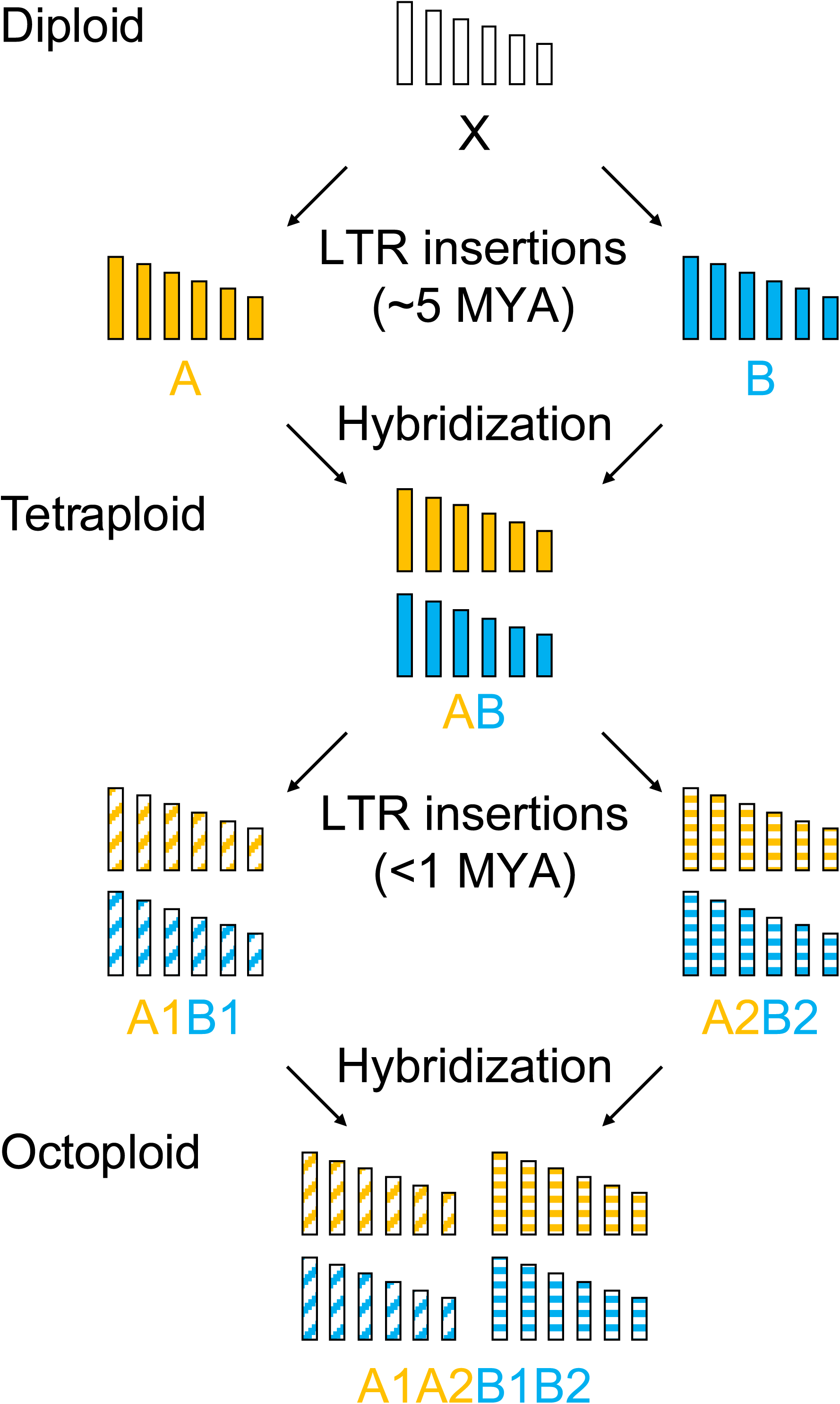
Proposed evolutionary model for the origin of octoploid *Oxalis corniculata*. The schematic illustrates a two-step hybridization and polyploidization process. Initially, two ancestral diploid lineages (A and B) diverged from a common ancestor (X) approximately 5 million years ago (MYA), as estimated by subgenome-specific LTR insertion times. These lineages hybridized to form an allotetraploid intermediate (AB). Subsequently, a second round of hybridization between differentiated tetraploid lineages (A1B1 and A2B2) occurred less than 1 MYA, resulting in the establishment of the extant octoploid *O. corniculata* (A1A2B1B2). Vertical bars represent the six chromosomes of each subgenome, with different colors and patterns indicating subgenome-specific sequence signatures acquired during their independent evolution.

Large-scale, spatially replicated sampling is essential for studying urban evolution, a goal we achieved through the *Minna de Katabami* ("Oxalis, Together") project, where more than 37 participants contributed 1,784 samples nationwide (Supplementary Table S2). While the morphological similarity between *Oxalis* species led to a 20% misidentification rate (323/1,612), this was not unexpected given that accurate species identification of *Oxalis* is challenging even for specialists (Moura et al., 2020). This is a recognized hurdle in large-scale citizen science projects (Roman et al., 2017). Furthermore, 172 samples were excluded due to low DNA quality or quantity, likely because the optimal condition of plant materials for DNA extraction is difficult for non-experts to judge. We effectively addressed these issues using genomic filtering (Figure 5), but these challenges highlight the importance of active engagement from researchers to bridge the gap between public participation and technical requirements. By increasing such interventions, scientists can more effectively share the fascination of DNA research with the public, evolving citizen science into a deeper opportunity for mutual learning. Despite these logistical hurdles, this distributed sampling was crucial for detecting nationwide repeatability; we found that while red and green forms occur widely across Japan, the genetic architecture was not identical, as Groups II and III showed a GWAS peak consistent with the mapped *MYB* locus while Group I did not (Figure 3, Supplementary Figure S7). This suggests that phenotypic evolution can be repeatable even when the underlying genetic basis is contingent on population history and standing genetic variation (Rosenblum et al., 2014; Therkildsen et al., 2019; Rêgo et al., 2025).

Our study advances urban evolutionary genomics, yet several limitations remain. While we identified a strong correlation between landscape and allele frequency, future work should include transplant experiments and temporal monitoring to estimate selection coefficients. Furthermore, functional validation of the candidate *MYB* variant will be necessary to clarify the genotype–phenotype correlation. Finally, identifying the alternative genetic mechanisms in Group I may reveal further cases of convergent evolution. Through such integrative and collaborative approaches, urban landscapes serve as powerful systems for forecasting evolutionary responses to human-driven environmental change, while simultaneously enhancing public engagement and education regarding evolutionary theory.

In conclusion, we demonstrate that urban landscape structure can shape adaptive genetic variation, resolvable at the level of specific loci. By combining high-quality polyploid genome sequences with the power of citizen science, our study highlights how everyday urban environments provide a unique laboratory for understanding evolution in a rapidly changing world.

## Materials and Methods

### Plant materials and anthocyanin content measurement

For genome sequencing and analysis, red- and green-leaved lines of *O. corniculata* collected from Tokyo, Japan, were used. These lines were previously described in our earlier study (Fukano et al., 2023b). For linkage analysis, an F2 mapping population was developed by self-crossing an F1 plant derived from a cross between the red- and green-leaved lines. In addition, we used a natural population of *O. corniculata* plants collected from 75 regions across Japan (Supplementary Table S2), with the cooperation of project participants. Anthocyanin content in the leaves was measured using an ACM-200 plus Anthocyanin Content Meter (Opti-Sciences, Hudson, NH, USA).

### Chromosome observation

Pollen from 1-mm buds was used for the observation of mitotic chromosomes. Inflorescences containing 1-mm buds were pretreated with 2 mM 8-hydroxyquinoline at room temperature for 3 hours and fixed in a 3:1 (v/v) ethanol–acetic acid solution at room temperature for 2 days. The fixed samples were stored in 70% ethanol at 4°C until use. Chromosomes were observed using the squash method described by Wang et al. (2015). Chromosomes were mounted with 5 μg/mL DAPI (Nacalai Tesque, 19178-91) in VECTASHIELD (Vector Laboratories, H-1000) and observed using OLYMPUS BX-53 fluorescence microscope.

### Genome sequencing and assembly

Genome DNA was extracted from young leaves using Genomic-tip 500/G (QIAGEN, 10262). To estimate the genome size of the red- and green-leaved lines, a short-read sequencing library was prepared using the xGen DNA EZ Library Prep Kit (Integrated DNA Technologies, Coralville, IA, USA) and converted into a DNA nanoball sequencing library with an MGI Easy Universal Library Conversion Kit (MGI Tech, Shenzhen, China). The library was sequenced on a DNBSEQ G400RS (MGI Tech) instrument in paired-end, 101-bp mode. The obtained reads were used to estimate genome size using a *k*-mer distribution analysis of Jellyfish (*k*-mer size = 17) (Marçais and Kingsford, 2011).

For the genome assembly, genome DNA was sheared to an average fragment size of 40 kb using a Megaruptor 2 (Diagenode, Liege, Belgium) in Large Fragment Hydropore mode. The sheared DNA was used for HiFi SMRTbell library preparation with the SMRTbell Express Template Prep Kit 2.0 (PacBio, Menlo Park, CA, USA). The obtained DNA libraries were fractionated with a BluePippin (Sage Science, Beverly, MA, USA) to eliminate fragments shorter than 20 kb. The fractionated DNA libraries were sequenced on a SMRT Cell 8M using the Sequel II system (PacBio). In parallel, a high-throughput chromosome conformation capture library was prepared using the Omni-C Proximity Ligation Assay (Dovetail Genomics, Scotts Valley, CA, USA) and sequenced on a DNBSEQ G400RS (MGI Tech) instrument in paired-end, 151-bp mode.

The HiFi reads and Omni-C reads were assembled using Hifiasm (version 0.19.9) (Cheng *et al*., 2021) in Hi-C integration mode. The --hg-size 1g and --hom-cov 33 options were used to obtain accurate homozygous read coverage. Haplotigs and overlaps in the assembled contigs were removed using Purge_Dups (version 1.2.6) (Guan *et al*., 2020) with the cutoffs assigned using calcuts -l 5 -m 20 -u 75. Contigs predicted to be split chromosomes were concatenated manually using HiFi reads that aligned contiguously to the ends of the two contigs by self-alignment in the green individual. Because many contigs in the assembly of the red individual were predicted to be split chromosomes, these contigs were scaffolded using RaGOO (version 1.1) with default parameters, using the assembly of the green individual as a reference (Alonge et al., 2019). Assembly completeness was evaluated using Benchmarking Universal Single-Copy Orthologs (BUSCO, version 5.5.0) with the -m genome, -l embryophyta_odb10, and --augustus options (Simão *et al*., 2015). Telomere sequences were detected using the search command of the Telomere Identification toolKit (tidk, version 0.2.31) with the -s TTTAGGG and --window 1000 options (Brown et al., 2025).

The organelle genomes were assembled from the HiFi reads using Oatk (Zhou *et al*., 2025) with a syncmer coverage threshold (-c) of 200. For the gene prediction of chloroplast genomes, reported genes from *O. corniculata* (GenBank accession number NC_051971.1) were aligned to the chloroplast genome assemblies from this study using LiftOn (Chao *et al*., 2025). Protein-coding, transfer RNA, and ribosomal RNA genes, as well as inverted repeats in the chloroplast genomes, were then predicted using CPGAVAS2 (Shi *et al*., 2019). For the gene prediction of mitochondrial genomes, protein-coding, transfer RNA, and ribosomal RNA genes were predicted using PMGA (Li *et al*., 2025). Gene positions on the organelle genomes were manually curated and corrected. The chloroplast and the mitochondrial genome annotations were visualized using OGDRAW (version 1.3.1) (Greiner et al., 2019).

### Prediction of protein-coding genes

Total RNA was extracted from leaves using the FavorPrep Tissue Total RNA Extraction Mini Kit (Favorgen, Ping-Tung, Taiwan). RNA-seq libraries were constructed using the Kinnex Full-Length RNA Kit and sequenced on the Revio system (PacBio). The obtained full-length cDNA sequences were processed using Iso-Seq to generate non-redundant transcript sequences, and open reading frames (ORFs) within the transcripts were predicted using TransDecoder (Haas, B.J.; https://github.com/TransDecoder/TransDecoder). The resulting deduced amino-acid sequences were used as a training gene set for (Brůna *et al*., 2023) to predict protein-coding genes in the genome assemblies.

### Repetitive sequence analysis and subgenome identification

Repetitive sequences in the assembly were identified using RepeatMasker (https://www.repeatmasker.org) based on repeat sequences registered in Repbase and a de novo repeat library built with RepeatModeler (https://www.repeatmasker.org).

The octoploid genome of *O. corniculata* was classified into subgenomes by examining the abundance of specific transposable elements (TEs) on specific chromosomes. Combinations of the top 12 chromosomes with the highest number of TEs per 1 Mbp were extracted for each TE, and the number of occurrences of these chromosome combinations was counted. The most frequently appearing combination of chromosomes was estimated to be a subgenome. Subphaser (Jia *et al*., 2022) was used to validate the subgenome classifications and to estimate the divergent time of the subgenomes.

### GWAS and genetic diversity analysis

Genome DNA was extracted from the leaves of the mapping and natural populations using the sbeadex DNA Extraction Kit on the oKtopure system (LGC, United Kingdom). The extracted genomic DNA was digested with PstI and MspI restriction endonucleases, subjected to ddRAD-Seq library preparation (Shirasawa et al., 2016), and sequenced on a DNBSEQ G400 instrument (MGI Tech). Subsequently, after removing adapter sequences (AGATCGGAAGAGC) using fastx_clipper in the FASTX-Toolkit (version 0.0.14; http://hannonlab.cshl.edu/fastx_toolkit) and trimming low-quality reads (quality score < 10) with PRINSEQ (Schmieder and Edwards, 2011), the ddRAD-Seq reads were aligned to the chromosome-level pseudomolecule sequences to detect sequence variants. High-confidence biallelic SNPs were selected using VCFtools (Danecek *et al*., 2011) with the following filtering conditions: read depth ≥ 5, SNP quality = 999, minor allele frequency ≥ 0.25 (for the mapping population) or ≥ 0.05 (for the natural population), and proportion of missing data < 20%. A phylogenetic tree, based on 100 bootstrap replicates, was constructed using SNPhylo (Lee et al., 2014) and visualized with iTOL (Letunic and Bork, 2021). Association mapping for quantitative traits was performed using a general linear model (for the mapping population and the subpopulation of the natural population) and a mixed linear model (for the entire natural population) implemented in TASSEL version 5.0 (Bradbury *et al*., 2007). Manhattan plots were drawn using qqman (Turner, 2018).

### RT-PCR and Sanger sequencing

Total RNA was extracted from leaves using the FavorPrep Plant Total RNA Extraction Mini Kit (Favorgen, Ping Tung, Taiwan) or FavorPrep Plant Total RNA Mini Kit (for woody plant) (Favorgen). cDNA was synthesized with gene-specific primers (5’-TGCAAAAGTAATACTTTAATCGATCGCT-3’ and 5’-ACAGTCGTACTCACGAGAACTG-3’) and electrophoresed on 1% agarose gels. Gene-specific DNA fragments were excised from the gels using a Gel/PCR Extraction Kit (FastGene, Tokyo, Japan). The nucleotide sequences of the DNA fragments were determined using the BigDye Terminator Cycle Sequencing Kit (Applied Biosystems, Waltham, MA, USA) and an ABI 3730xl DNA Analyzer (Applied Biosystems). The data were analyzed using Unipro UGENE (Okonechnikov et al., 2012), and primers were designed using Primer3 (Koressaar and Remm, 2007).

### Fragment analysis

PCR was performed using 1 ng of genomic DNA in a 5 μL reaction mix containing 1× NH_4_ buffer, 2 mM MgClL, 0.125 U BIOTAQ™ DNA polymerase (BIOLINE), 2 mM dNTPs, 0.25 μM of a forward primer (5’-CCACTTTCAACGAGCTGATGGCCTCGGACCTTCAAGCTTAA-3’), 0.5 μM of a reverse primer (5’-CTGATGATGATTAACAAAATCATCAGGATCATAT-3’), and 0.125 μM of a fluorescent labeled primer (5’-CCACTTTCAACGAGCTGATG-3’). Thermal cycling conditions were as follows: an initial denaturation at 94°C for 2 min; 40 cycles of denaturation at 94°C for 30 s, annealing at 55°C for 30 s, and extension at 72°C for 60 s; and a final extension at 72°C for 3 min. PCR products were separated using a 3730 DNA Analyzer (Applied Biosystems), and the data were analyzed using GeneMarker software (SoftGenetics, State College, PA, USA).

## Acknowledgements

We are grateful to the participants of the ’Minna de Katabami’ Project for their contribution to sample collection across Japan: K. Sugimoto (Tsukuba); K. Nashima (Fujisawa); Hisanaga and members of Igusa High School, Tokyo (Nerima); S. Utsumi (Sapporo); A. Ichimura, N. Suzuki, R. Sakurai, and members of Chiba Eiwa High School (Yachiyo); K. Kitagawa (Goto); members of the Biological Research Club at Tennoji High School, Osaka (Osaka); K. Fujii and H. Tabata (Bunkyo); T. Hanada, M. Maeda, M. Sakamoto, A. Shinohara, S. Hanada, K. Nakayama, and H. Miyazaki (Kitakyushu); members of the Biology Club at Rikkyo Niiza High School, Saitama (Niiza); S. Okamoto (Konan); Y. Agawa (Kushimoto); H. Sassa (Minamidaito); R. Sato (Fukuoka and Hita); R. Morishita (Nagano); K. Murayama (Sapporo); members of Matsudo Mutsumi High School, Chiba (Matsudo); N. Okada, N. Iwatani, K. Yoshida, Y. Sugimura, M. Fujibuchi, K. Shimizu, and M. Seki (Fujieda); T. Toda (Nagoya); K. Hori (Kochi); and other anonymous contributors. We thank Y. Kishida, C. Minami, K. Ozawa, H. Tsuruoka, and A. Watanabe (Kazusa DNA Research Institute) for their technical assistance.

## Funding

This work was supported by JSPS KAKENHI (Grant Numbers 16H06279 [PAGS], 21H02559, 22H05172, 22H05181, and 24K02096) and the Kazusa DNA Research Institute Foundation.

## Data availability

Raw sequence reads and assembled sequences are available at DDBJ (BioProject accession number PRJDB39843) and Kazusa Genome Atlas (https://genome.kazusa.or.jp).

## Supplementary Information

**Supplementary Table S1.** Statistics of the initial contig assemblies for the red- and green-leaved *Oxalis corniculata* lines.

**Supplementary Table S2.** List of *Oxalis corniculata* samples collected from across Japan.

**Supplementary Figure S1** Chromosomes of *Oxalis corniculata*.

Meiotic chromosomes for (A) the red-leaved line and (B) the green-leaved line. Scale bar = 5 µm.

**Supplementary Figure S2** Genome size estimation for the red- and green-leaved lines.

*K*-mer distribution (*k* = 17) showing the frequency of distinct *k*-mers against their multiplicity values.

**Supplementary Figure S3** Subgenome-specific repetitive sequences.

Counts of repetitive sequences enriched in (A) subgenome A, (B) subgenome B, and (C) subgenome 1.

**Supplementary Figure S4** Subgenome phasing based on repetitive *k*-mers.

A–D: Chromosomal characteristics for the red-leaved line (A: subgenomes A and B; B: subgenomes 1 and 2) and the green-leaved line (C: subgenomes A and B; D: subgenomes 1 and 2). Concentric rings from outer to inner represent:

1. Subgenome assignments determined by a *k*-means clustering algorithm.
2. Significant enrichment of subgenome-specific *k*-mers.
3. Normalized proportions of subgenome-specific *k*-mers.

(4–5) Density distribution of each subgenome-specific *k*-mer set.

1. Density distribution of subgenome-specific LTRs and other LTRs.
2. Homoeologous blocks across each homoeologous chromosome set.

E–H: Insertion times of subgenome-specific LTRs for the red-leaved line (E: subgenomes A and B; F: subgenomes 1 and 2) and the green-leaved line (G: subgenomes A and B; H: subgenomes 1 and 2).

**Supplementary Figure S5** Organelle genomes of *Oxalis corniculata*.

Chloroplast genomes for (A) the red-leaved line and (B) the green-leaved line. Mitochondrial genomes (represented by two circular contigs) for (C, D) the red-leaved line and (E, F) the green-leaved line.

**Supplementary Figure S6** Histograms of Anthocyanin Color Index (ACI) representing leaf color variation. Frequency distribution of ACI values for (A) the F2 mapping population and (B) the natural population collected from across Japan.

**Supplementary Figure S7** SNPs associated with leaf color variation identified by GWAS.

Manhattan plots showing the distribution of SNPs associated with leaf color variation across the genome. The horizontal red lines indicate the Bonferroni-corrected significance threshold (α = 1%). Results are shown for (A) the F2 mapping population using the green-leaved line assembly as a reference, and (B–D) Groups I, II, and III of the natural population collected from across Japan.

## Notes

### Competing Interest Statement

The authors have declared no competing interest.

### Summary of Updates

Corrected author names and updated ORCID information.

