## Supplementary Figures for "Genomic basis of rapid urban evolution revealed by the subgenome-resolved genome of octoploid *Oxalis corniculata*"

**Supplementary Information**

**Supplementary Table S1** Statistics of the initial contig assemblies for the red- and green-leaved *Oxalis corniculata* lines.

**Supplementary Table S2** List of *Oxalis corniculata* samples collected from across Japan.

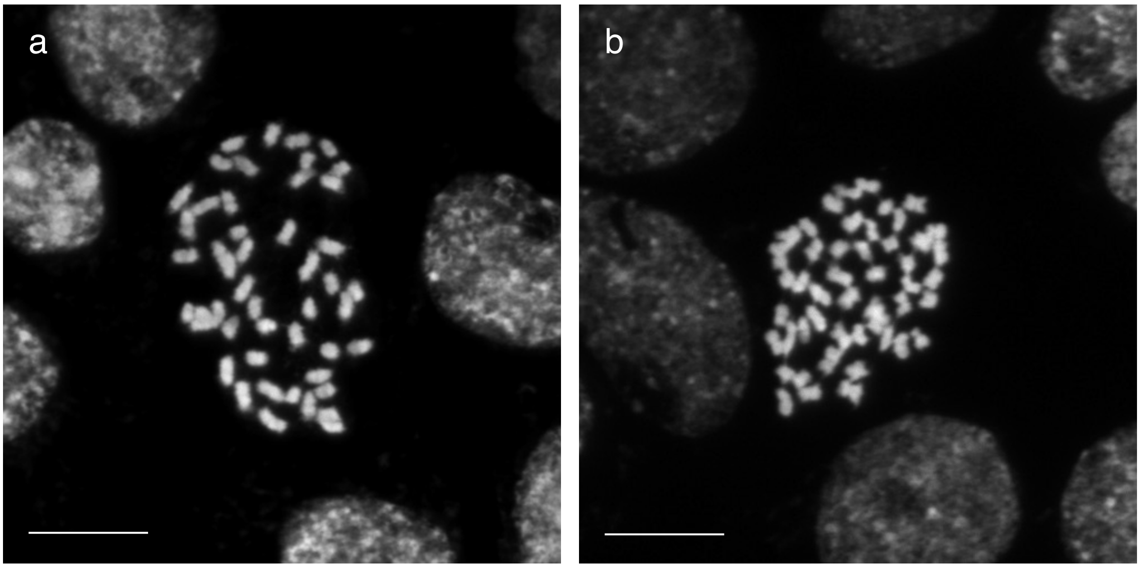


**Supplementary Figure S1** Chromosomes of *Oxalis corniculata*.

Meiotic chromosomes for (A) the red-leaved line and (B) the green-leaved line. Scale bar = 5 µm.

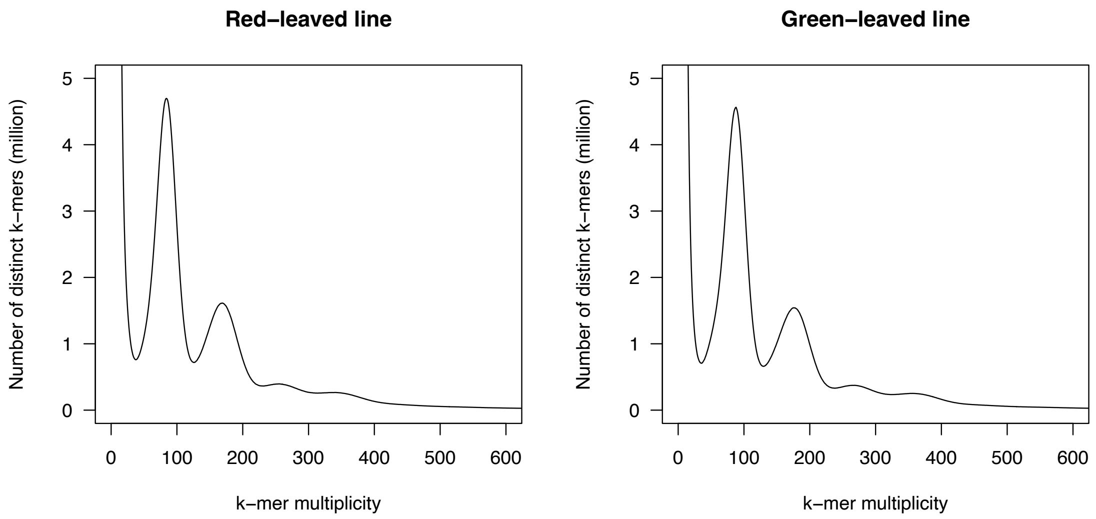


**Supplementary Figure S2** Genome size estimation for the red- and green-leaved lines. *K*-mer distribution ($k = 17$) showing the frequency of distinct *k*-mers against their multiplicity values.

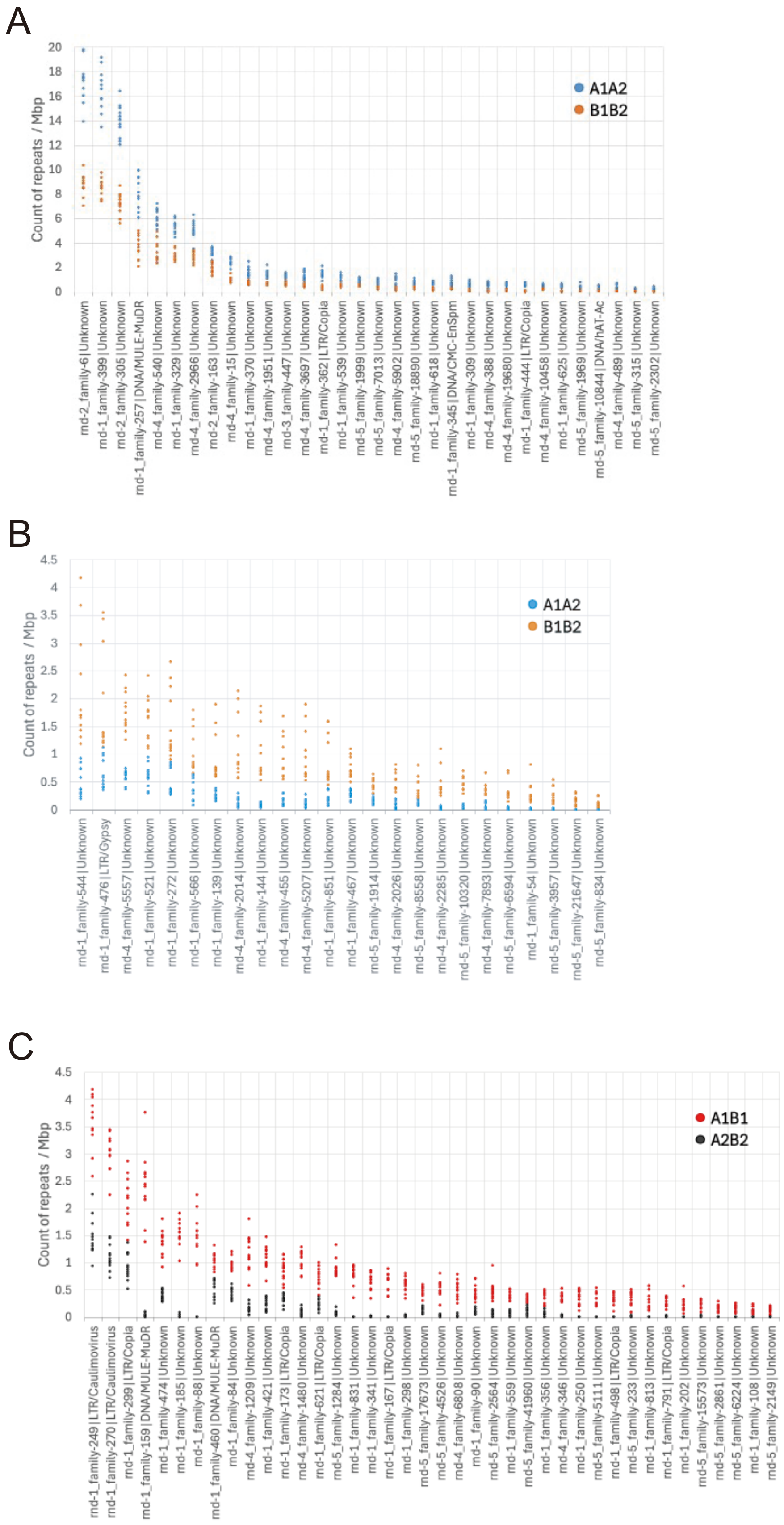


**Supplementary Figure S3** Subgenome-specific repetitive sequences.

Counts of repetitive sequences enriched in (A) subgenome A, (B) subgenome B, and (C) subgenome 1.

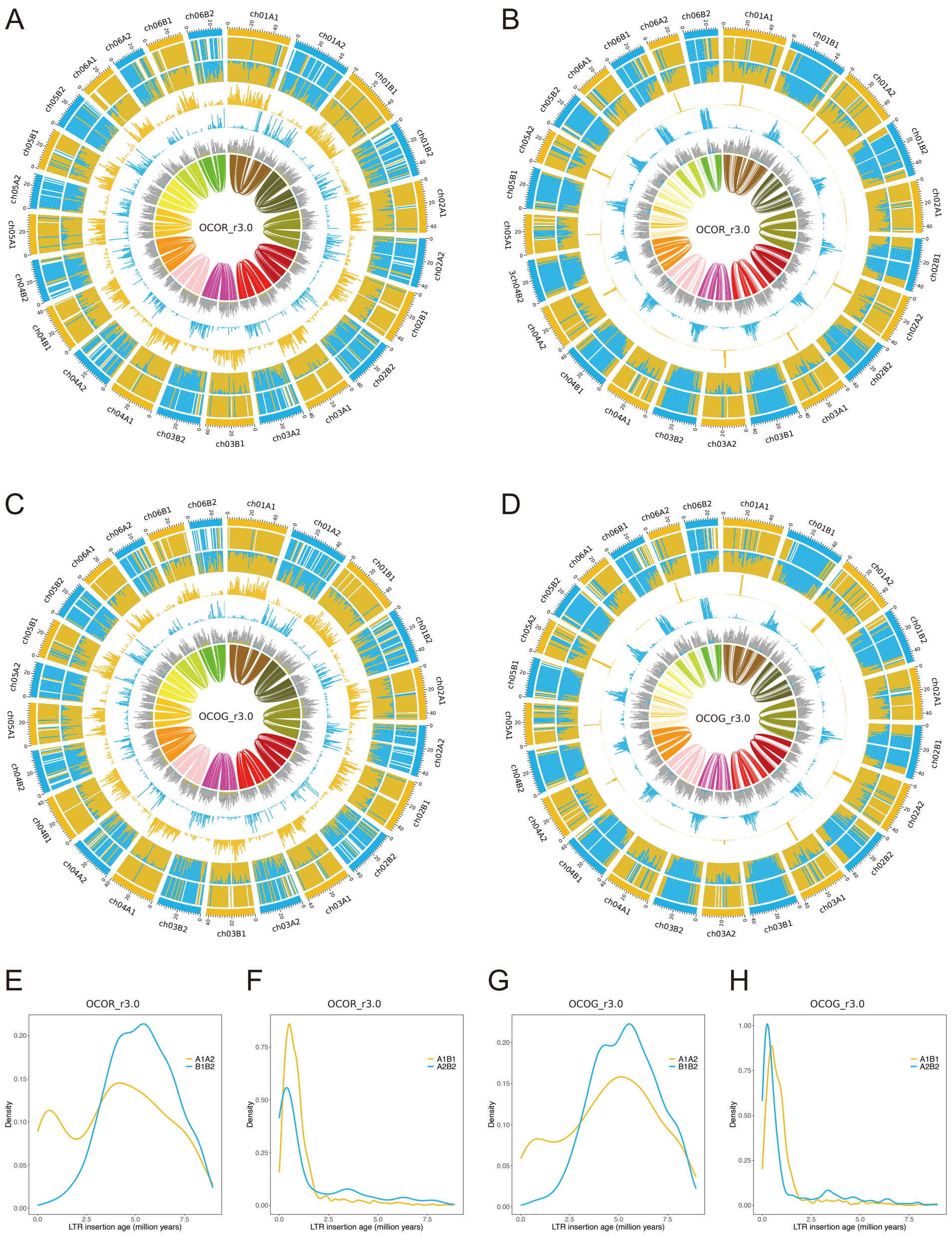


**Supplementary Figure S4** Subgenome phasing based on repetitive *k*-mers.

A–D: Chromosomal characteristics for the red-leaved line (A: subgenomes A and B; B: subgenomes 1 and 2) and the green-leaved line (C: subgenomes A and B; D: subgenomes 1 and 2). Concentric rings from outer to inner represent:

(1) Subgenome assignments determined by a *k*-means clustering algorithm.

(2) Significant enrichment of subgenome-specific *k*-mers.

(3) Normalized proportions of subgenome-specific *k*-mers.

(4–5) Density distribution of each subgenome-specific *k*-mer set.

(6) Density distribution of subgenome-specific LTRs and other LTRs.

(7) Homoeologous blocks across each homoeologous chromosome set.

E–H: Insertion times of subgenome-specific LTRs for the red-leaved line (E: subgenomes A and B; F: subgenomes 1 and 2) and the green-leaved line (G: subgenomes A and B; H: subgenomes 1 and 2).

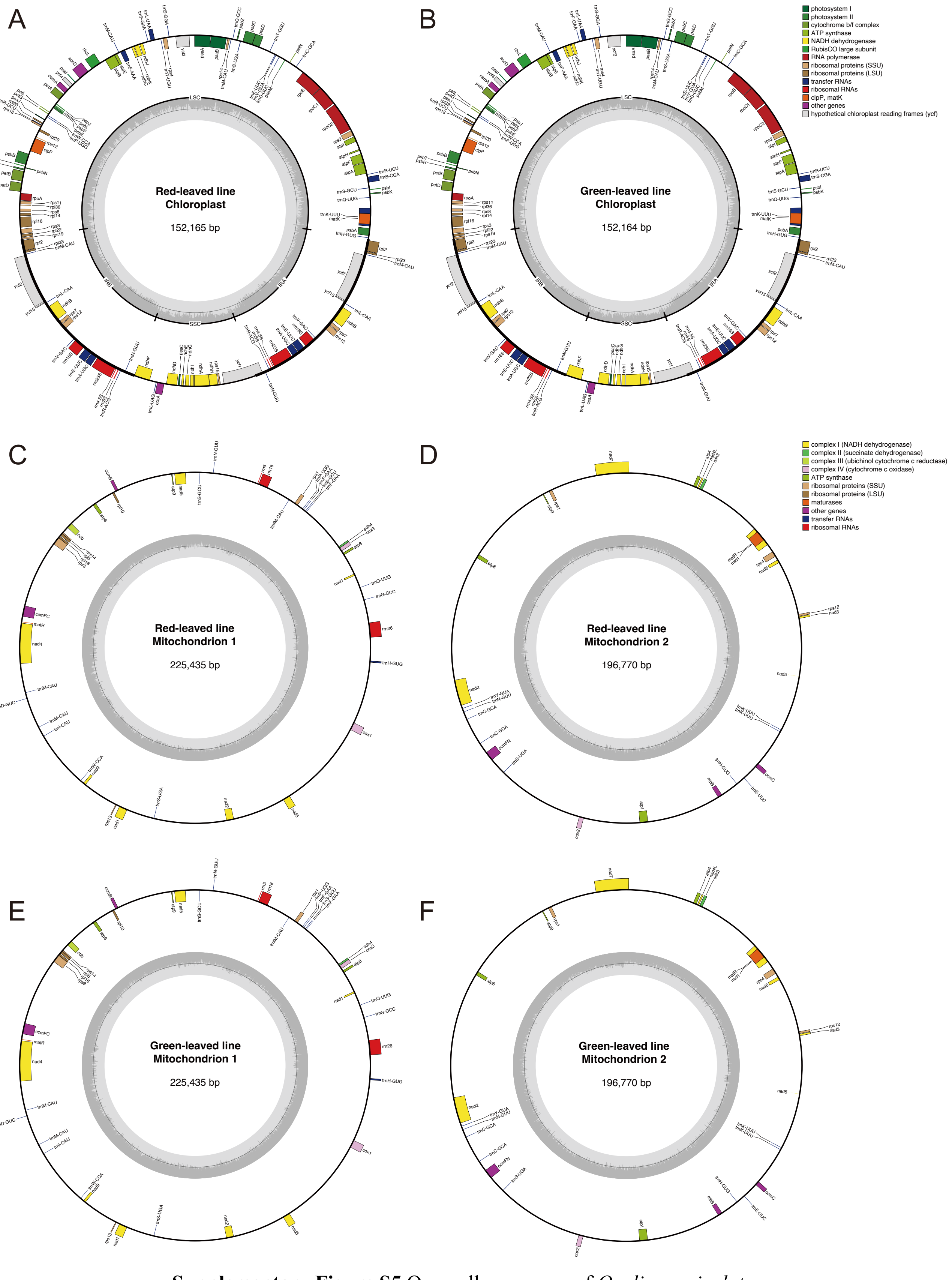


**Supplementary Figure S5** Organelle genomes of *Oxalis corniculata*.

Chloroplast genomes for (A) the red-leaved line and (B) the green-leaved line. Mitochondrial genomes (represented by two circular contigs) for (C, D) the red-leaved line and (E, F) the green-leaved line.

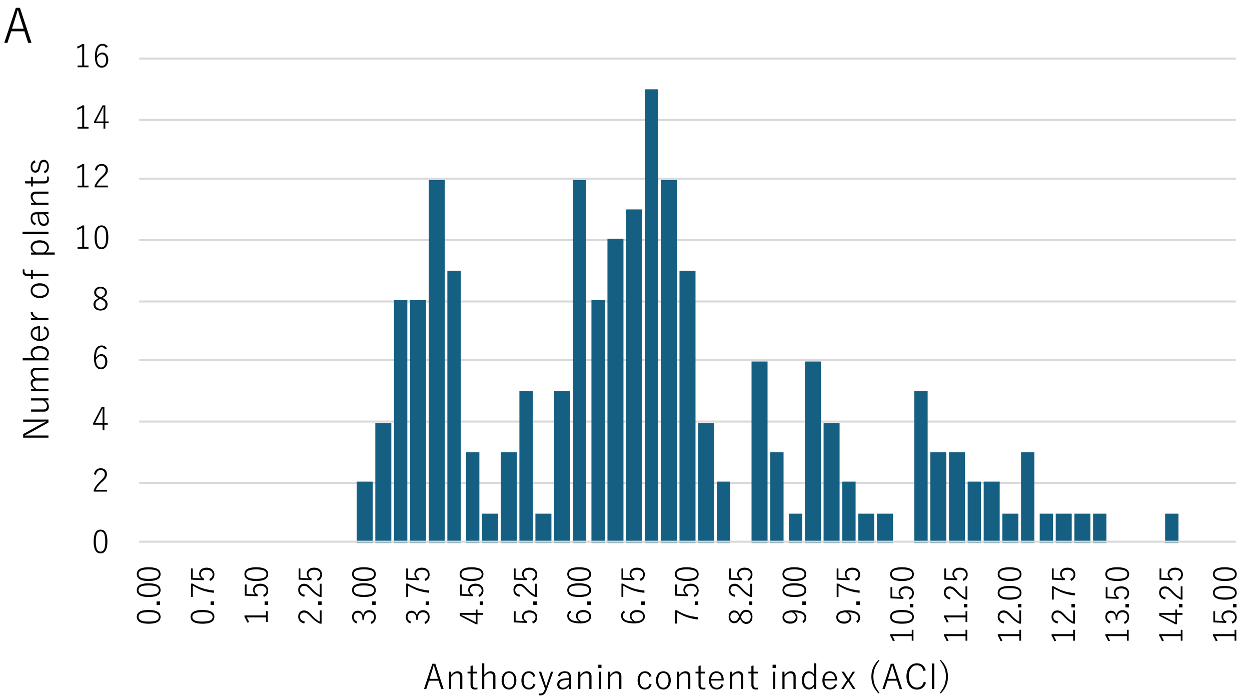


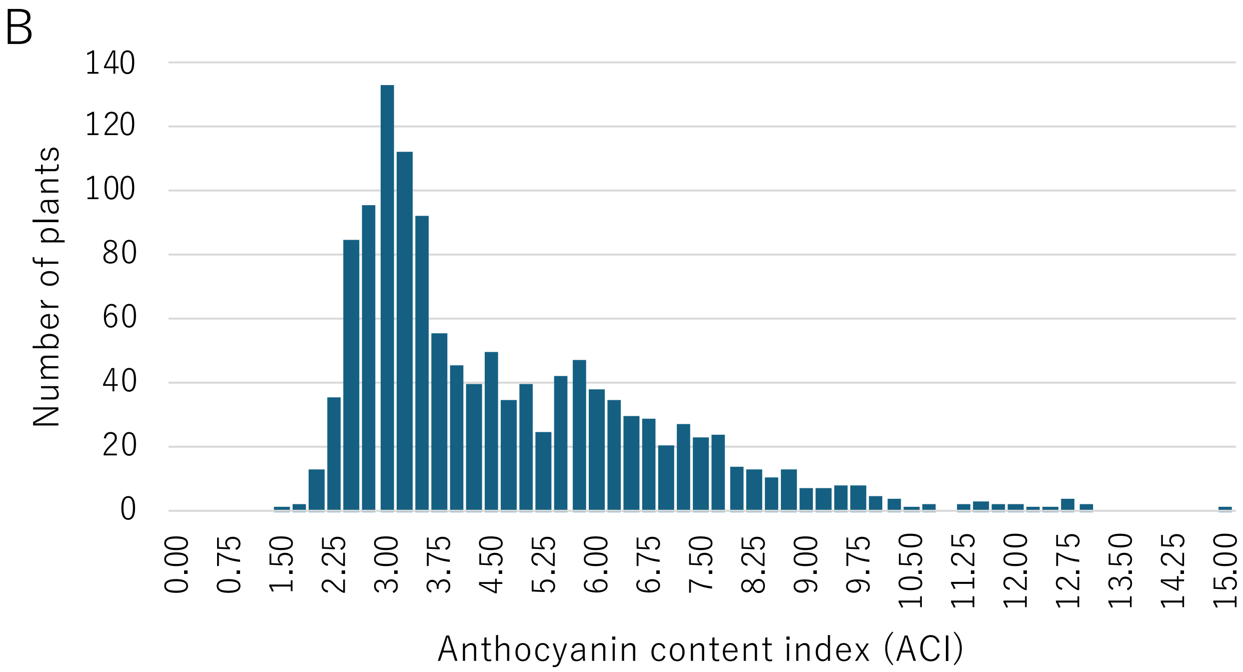


**Supplementary Figure S6** Histograms of Anthocyanin Color Index (ACI) representing leaf color variation. Frequency distribution of ACI values for (A) the F2 mapping population and (B) the natural population collected from across Japan.

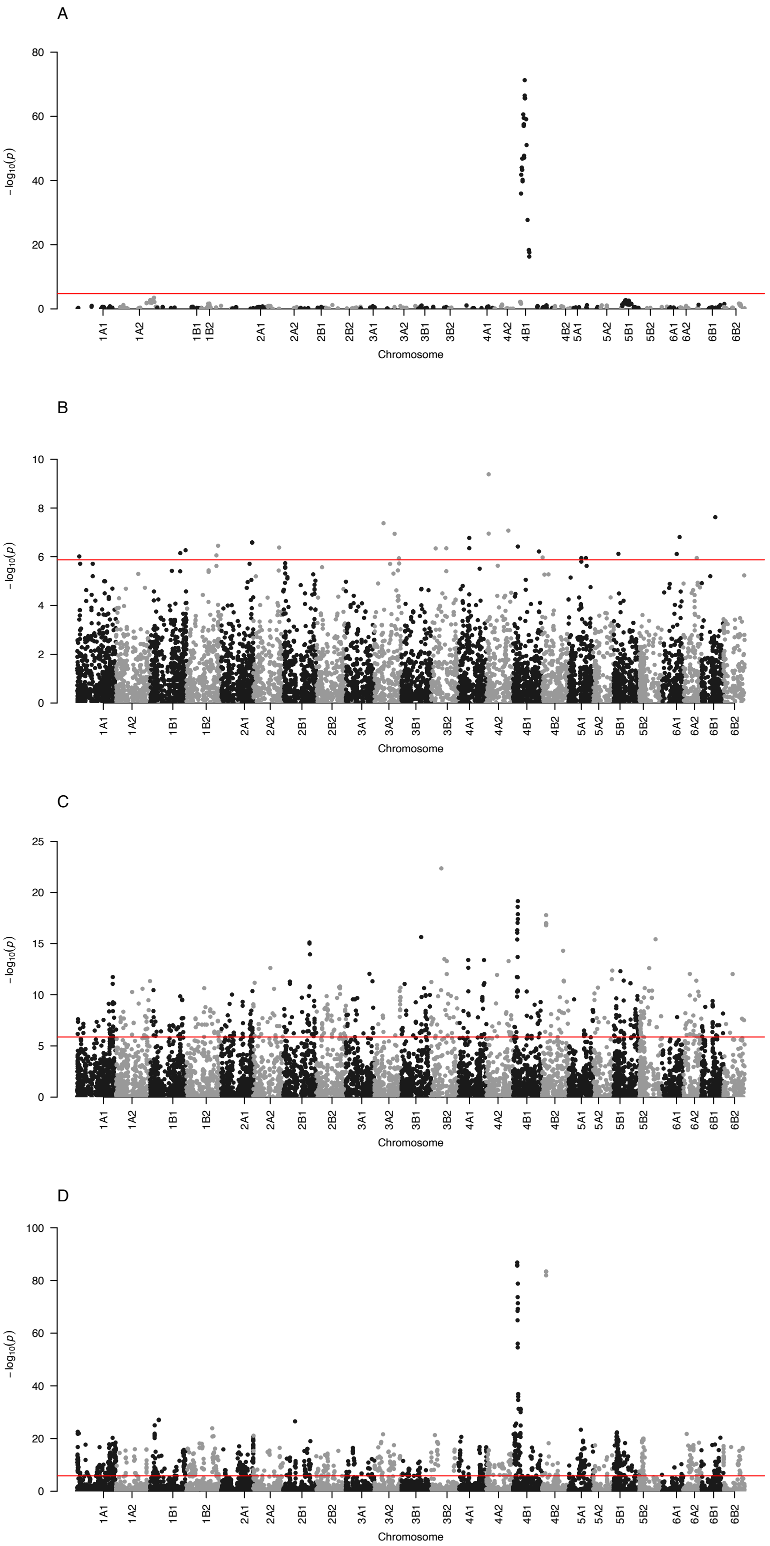


**Supplementary Figure S7** SNPs associated with leaf color variation identified by GWAS.

Manhattan plots showing the distribution of SNPs associated with leaf color variation across the genome. The horizontal red lines indicate the Bonferroni-corrected significance threshold ($\alpha = 1\%$). Results are shown for (A) the F2 mapping population using the green-leaved line assembly as a reference, and (B–D) Groups I, II, and III of the natural population collected from across Japan.
